# Retrieval of whole human genome clinical variant information through search motors

**DOI:** 10.1101/600098

**Authors:** Li Yin, Julia di Iulio, Sebastien Lelong, Chunlei Wu, Amalio Telenti

## Abstract

The interpretation of variation in the human genome constitutes one of the most pressing challenges in biomedicine. There are many academic and proprietary resources that provide annotation, interpretation, scoring and knowledge of genetic variants under multiple access modalities. For these resources, information is available through portals, wikis, APIs, curated content and databases, and pipelines. Here we explore the use of search motors to provide facilitated access to the main sources of information that are used by clinical geneticists and researchers. We also support the browsing experience with natural language processing and automated summaries of available information. The resulting tool, ai-OMNI.com, is intrinsically flexible, expandable and intuitive, thus providing a different experience for querying the human genome for the consequences of variation.

## Background & Summary

Genomic information is challenging because the recent changes in technology – from genotyping arrays to exome and to whole genome sequencing – are uncovering massive amounts of uncharacterized genetic variants. Recent efforts such as those of TopMed, gnomAD or from industry have so far identified over 800 million human variants^1-3^. Of these, there is clinical information for less than 250,000 variants (0.03%). The interpretation of common genetic variants is mostly established by statistical association in large population studies (such as GWAS – genome wide association studies), while the interpretation of rare variants is mainly performed through careful manual curation of clinical and functional information – generally in the setting of studies of a family pedigree. There are recommendations of the American College of Medical Genetics and Genomics for reporting of findings in clinical exome and genome sequencing^4^.

In clinical practice, the process of assessment of single or multiple genetic variants requires access to different sources of information tools and resources. This includes clinical information, genome information, prediction of deleteriousness of genetic variants, and literature resources. Many of these sources can be provided as “annotations” during the automated generation of genome sequence reports; however, for a limited number of variants, there is in-depth clinical work – in particular those labelled as “variants of unknown significance” or VUS^5^. This dedicated effort is today very time consuming by requiring access to disparate sources of information from portals, databases, wikis, APIs and curated resources and publications.

The aim of the present work is to develop a different approach to the frequently fragmented information on genomics that exploits the intuitive interface of a search motor. A web browser has the ability to load web pages rapidly and generate a ranked representation of answers to short queries that are easy to inspect and link to the original sources. We explored this tool as a more flexible approach to the increasing number of resources that are generally out of reach for the general public because they are only made available through data repositories, or from complex, not always user-friendly bioinformatic resources with a limited capacity for exploration. In addition, we explore the value of natural language processes to provide the general context (literature, text resources) on a variant of interest. **Table 1** presents a comparative of the new browser (https://ai-OMNI.com) with representative tools available to the community. OMNI allows users to view and interact with resources, tools and data repositories directly through a single search engine.

**Table 1.**
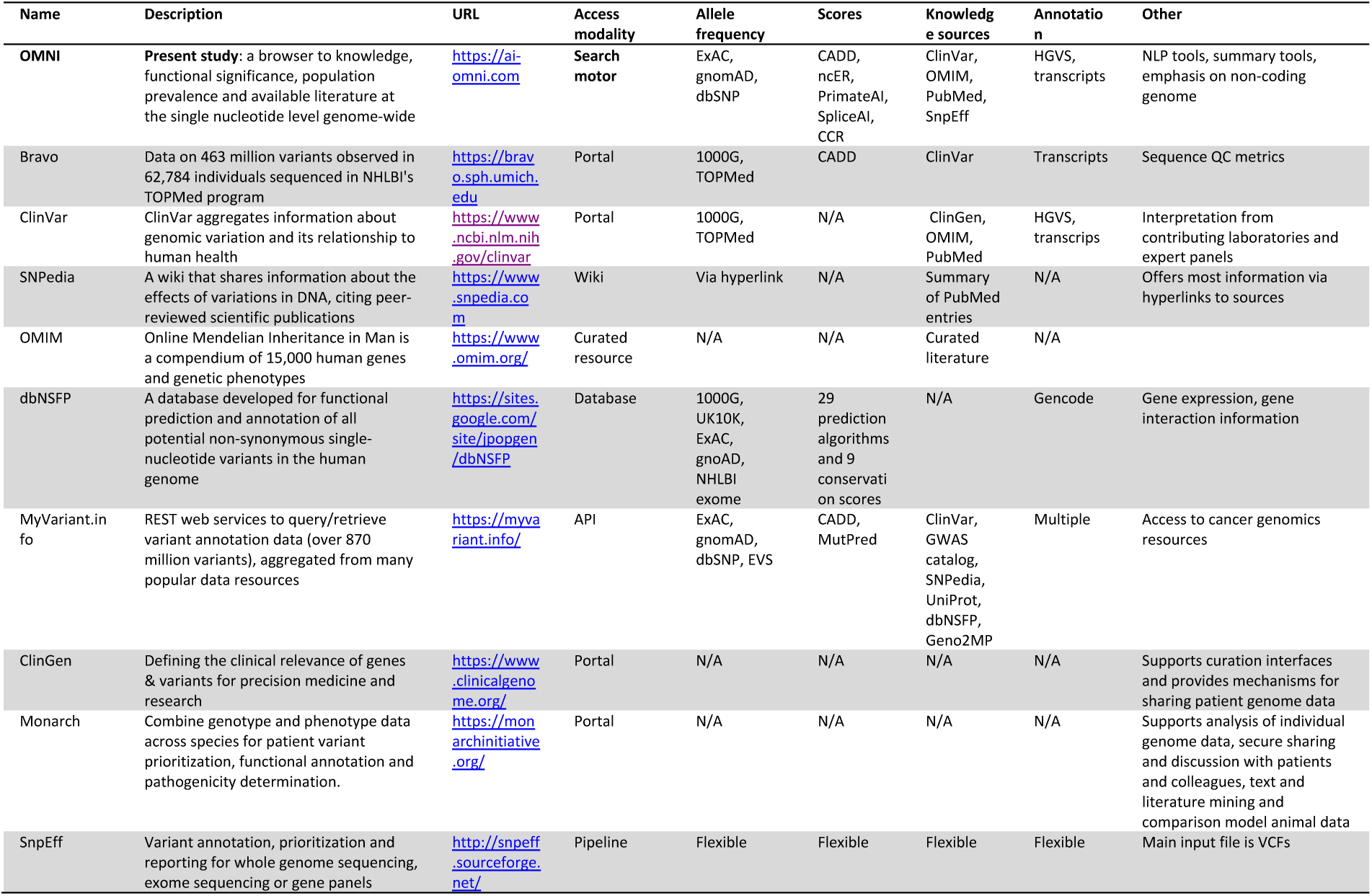
Comparison of the modalities of access and content of OMNI and a representative selection of other common resources in genomics. A portal usually focuses on hosting contents that are either self-produced or digested from other providers, and searchability is generally limited to the portal content. A wiki uses crowdsourcing that emphasizes the collaborative work by users to produce and modify content directly from the web browser. An application programming interface (API) is a set of functions and procedures allowing to communicate with data sources programatically. A software pipeline consists of a chain of code to achieve a task, such as genomic interpretation routine. A database is an organized storage of data and is often accessed via APIs. Content curation is the process of gathering information relevant to a particular topic by domain experts. Exome Aggregation Consortium (ExAC), Genome Aggregation Database (gnomAD), Combined Annotation Dependent Depletion (CADD), non-coding Regulatory Essential (ncER), Online Mendelian Inheritance in Man (OMIM), Human Genome Variation Society nomenclature (HGVS), Natural Language Processing (NLP), 1000 Genomes project (1000G), UK 10,000 genomes (UK10K), National Heart, Lung, and Blood Institute (NHLBI), Short Genetic Variations database (dbSNP), Exome Variant Server (EVS), Genome Wide Association Study (GWAS), Variant Call Format (VCF).

## Methods

The OMNI project consists of three major parts: the frontend app, the backend application programming interface (APIs) and the machine learning models. We created a web application, OMNI, using a modern JavaScript framework, React, to provide a responsive user interface (UI) that can search, fetch and visualize data from various data sources. We used asynchronous JavaScript to communicate to different APIs without freezing the UI. In addition to the existing public APIs, we also created dedicated APIs to perform Natural Language Processing (NLP), data indexing and secure data proxying. We created an Elasticsearch Index and exposed a REST API, which also included novel machine learning and deep learning scores: ncER, SpliceAI, PrimateAI and CCRs (**Figure 1**).

**Figure 1.**
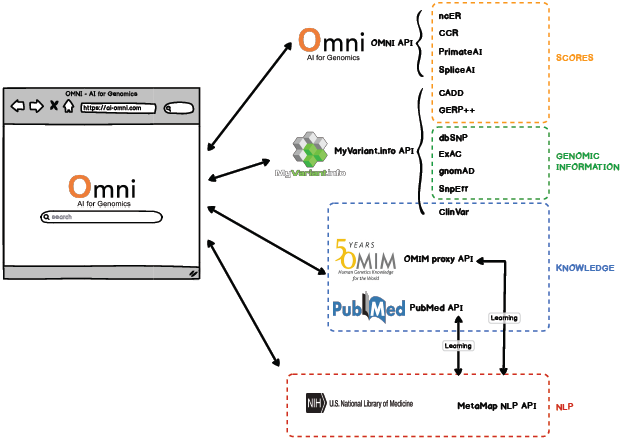
Content and sources of OMNI. The browser can search, fetch and visualize pathogenicity and deleteriousness scores, genomic detail and population frequencies at the single variant level, genome-wide. In addition, it provides access to existing primary knowledge and at summary level via natural language processing (NLP) of the text sources. All data remains unmodified and linked to the original source. Additional information on data sources is shown in **Suppl. Table S1**.

### Framework

The framework used for the frontend is React (https://reactjs.org/), which is one of the most popular and fast-growing frameworks for web development. It allows developers to create component-based applications. We used Redux (https://redux.js.org/) as the app state container and React-Router (https://reacttraining.com/react-router/) to manage app navigation. We also used Jest and Enzyme for testing, which will be covered in more details in the following section.

For the backend, we mainly used two Python frameworks: Flask and Tornado. Flask is used to build most of our app specific APIs, since it is lightweight to start with, and can add more functionalities on demand. APIs built with flask includes:

- An NLP REST API, which consumes literatures from public sources, such as PubMed, and perform NLP on them using a National Library of Medicine (NLM) licensed tool called MetaMap (https://metamap.nlm.nih.gov);
- A data proxy REST API, which serves limited data from Online Mendelian Inheritance in Man (OMIM).

These APIs are deployed with uWSGI (https://uwsgi-docs.readthedocs.io/en/latest/), which enables multiprocessing, and reverse-proxied with Nginx (https://www.nginx.com/).

We also build a nucleotide-level REST API under Tornado, a popular Python web framework for asynchronous networking, to serve the machine learning and deep learning scores on nucleotides. These scores are stored in MongoDB (https://www.mongodb.com/) and indexed using Elasticsearch (https://www.elastic.co/). This API is bootstrapped with BioThings SDK (http://docs.biothings.io/en/latest/), which allows us to build, index and serve large dataset conveniently. BioThings SDK is the same API framework used to power the high-performance MyVariant.info API^6^.

### Data sources

OMNI is currently exploring scores, genomic information and knowledge that are routinely used in clinical and research areas (**Figure 1**). In particular, the browser answers to queries by providing population allele frequency (dbSNP, gnomAD, ExAC)^1,3,7^, expected variant effects (SnpEff)^8^, a broadly used pathogenicity and conservation scores (CADD, Gerp++)^9,10^, and sources of information at variant level (ClinVar, OMIM, PubMed)^11,12^. In addition, the browser provides access to novel scores that are not broadly available and that are representative of a new family of machine learning genomic metrics for coding and non-coding regions^13-15^. The relevant links are available in **Supp. Table S1**.

## Technical Validation

OMNI does not modify or interpret the source data, scores or literature – it limits the scope of search to the representation and linking to the various resources (**Figure 2**). The only process that we currently test as novelty is the value of NLP to provide rapid inspection of the possible context of knowledge on a given genetic variant. Therefore, the technical validation herein refers to the testing of robustness and reliability of the technical platform.

**Figure 2.**
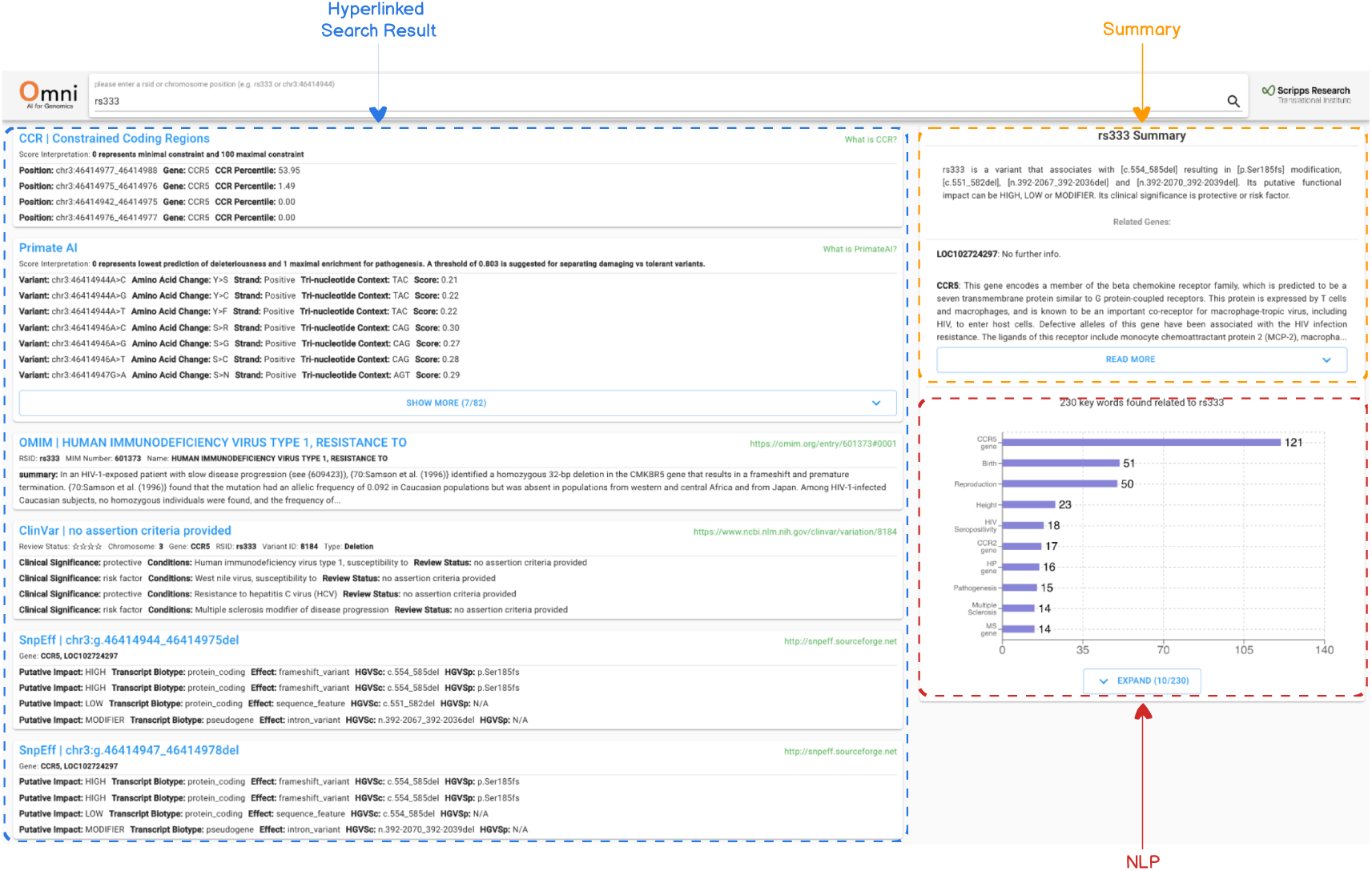
OMNI frontend. OMNI functions as a search motor that delivers original search engine results pages and also mines data available in databases. To support the user, OMNI also generates a summary statement on the variant and the associated gene. When publications prominently cite a unique variant in the abstract, or when the variant is included in Online Mendelian Inheritance in Man, the user can run a natural language process (NLP) analysis of those text resources.

For quality assurance, we used Jest (https://jestjs.io/) and Enzyme (https://airbnb.io/enzyme/) as the two major testing libraries. Jest provides developers a simulated environment of the browser and spy on the behaviours and interactions with the browser APIs to check if they work as expected. Enzyme provides a more accessible interface to the existing testing framework and makes the testing process more predictable.

In addition to the traditional unit testing and integration testing methodology, we adopted two underlying testing strategies called shallow rendering and snapshot testing. Since React app is component-based, we can test whether each component works individually (unit testing) and cooperates as expected (integration testing). Shallow rendering makes testing possible with minimum cost. It allows us to go only one level deep down the component tree without interfering with underlying components. It also provides the possibility to dive into deeper levels to perform integration testing. Snapshot testing, on the other hand, creates a full snapshot of the current component given its state, during the first run, and would compare against this snapshot in the future and warn about any inconsistency should it occur. Snapshot testing provides a much more detailed representation on the component quickly at the cost of being ignorant of the underlying logic. Combined with shallow rendering, snapshot testing can be extremely powerful to capture even the least significant change and lets the developer decide whether it is intentional or a potential bug.

OMNI also adopted continuous integration with CircleCI (https://circleci.com/) and auto-deployment with Heroku (https://dashboard.heroku.com/), so that whenever a new version has been pushed to the version control system, it will trigger CircleCI to run the tests first. After the tests has been passed, the app will be deployed automatically. It allows us to deliver new features constantly with confidence.

## Usage Notes

The API code and ncER dataset are available for download from our lab GitHub page (https://github.com/orgs/TelentiLab). Some resources are limited for sharing due to third-party copyright concern and may be available upon request from the respective owners.

## Discussion

Genome variant interpretation for the purpose of research or for clinical medicine is challenging. There are many available resources that provide datasets, scores, interpretation algorithms, guidelines panel recommendations, and specialized interest groups. The information is thus conveyed via portals, APIs, wikis, academic, institutional and proprietary support. OMNI aims at providing an agile interface that intersects many of the key resources that are used for clinical and research interpretation but that are, to a significant extent, dispersed throughout the various sources of information and knowledge. It does also experiment with automated extraction of salient information by NLP processes. However, OMNI applies a policy of no-modification or interpretation of the material as provided by the original sources – OMNI simply vehiculates the information and links to the original sites. In the future, OMNI will expand the number of modalities that are used in response to search queries (e.g., explicit information on the genomic element that contains a given variant, functional information emerging from genome-wide reporter and CRISPR-Cas9 screens), and also a dialog box to support user-initiated annotation of a given genetic variant. Finally, OMNI provides open-sourced APIs that facilitate access to resources that are difficult to query for the general user.

## Acknowledgements

Work of A.T. is supported by the Qualcomm Foundation and the NIH Center for Translational Science Award (CTSA, grant number UL1TR002550). Work of C.W. is supported by R01GM083924. We thank Xiaoqian Jiang and Kai Wang for useful input to this work.

## Author contributions

L.Y. developed the infrastructure and engineering of the site. J.dI. contributed genomic information and datasets. S.L. provided technical support for building a nucleotide-level REST API using BioThings SDK. C.W. provided technical guidance and access to the APIs of MyVariant.info. A.T. designed and supervised the study and wrote the paper with the help of co-authors.

## Competing interests

The authors declare no competing interests.

### Data availability

No primary data were used for this study.

---

**Supplementary Table S1.**
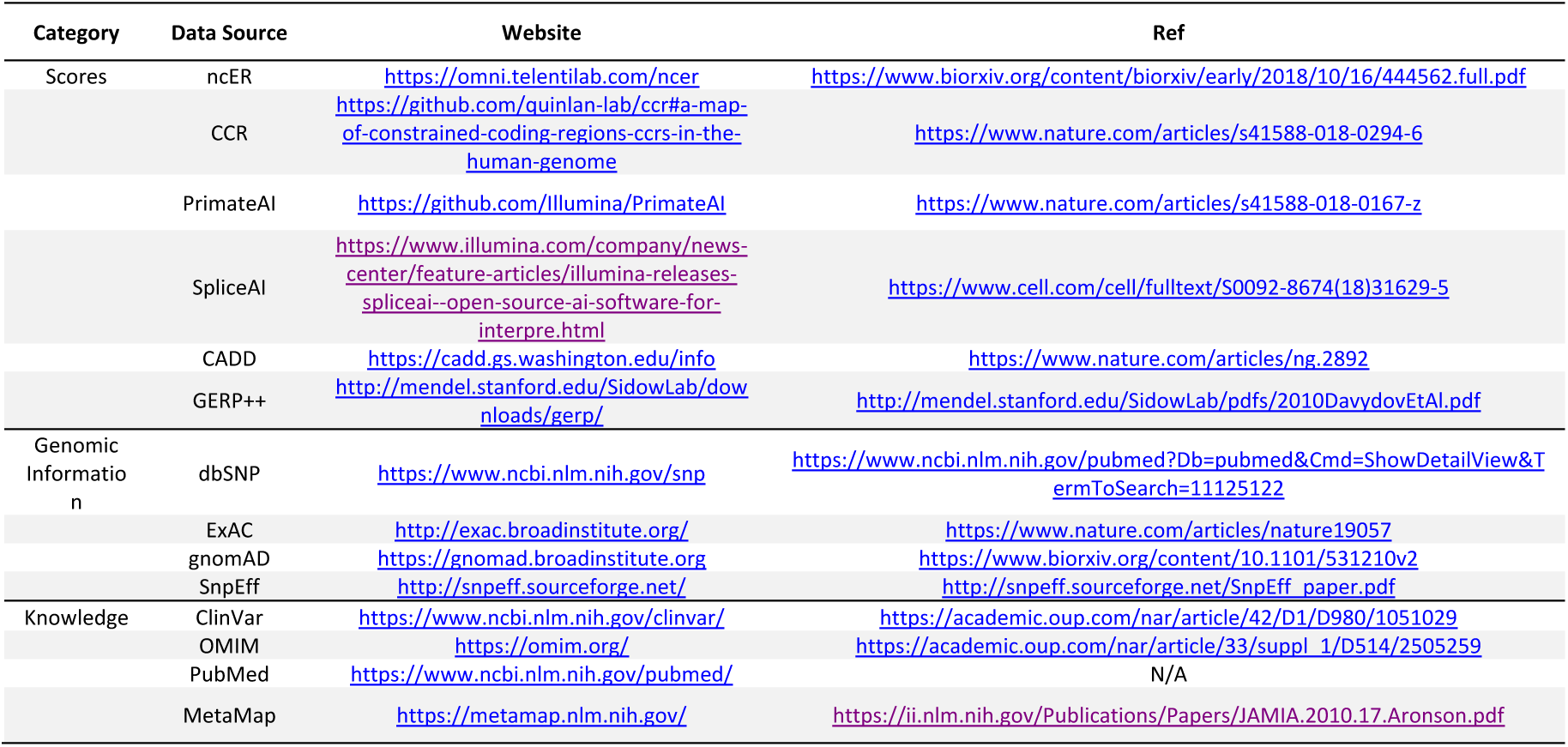
Data sources used in OMNI

